# Single-cell transcriptomics atlas of *Echinococcus granulosus* reveals germinative cell heterogeneity and candidate developmental key genes

**DOI:** 10.64898/2026.11.23.768474

**Authors:** Qian Wang, Jingwei Quan, Huaiyue Wang

**Author notes:** Corresponding author: Qian Wang. Authors contributed equally. **Authors’ contact information:** Qian Wang, Jingwei Quan, Huaiyue Wang.

## Abstract

Cystic echinococcosis is a zoonotic parasitic disease caused by infection with *Echinococcus granulosus*. Progressive cyst growth can compress host tissues and elicit immune responses. Characterizing the cellular composition and lineage differentiation of *E. granulosus* protoscoleces may provide insight into the regulation of parasite proliferation and development and facilitate the identification of potential intervention targets.

We generated a single-cell transcriptomic atlas of *E. granulosus* and identified six major cell populations: Germinative cell, Muscle, Neuron, Storage, Tegument,and Cilium. Germinative cells were heterogeneous: some retained an undifferentiated transcriptional profile, whereas others expressed genes associated with differentiated cell types, consistent with the initiation of lineage-associated programs. RNA velocity analysis suggested four potential trajectories, one involving transitions within the undifferentiated population that may contribute to self-renewal and homeostasis, and three showing trends toward distinct lineage-associated subpopulations. A *p53* homolog and threonine dehydrogenase *tdh* were identified as candidate developmental key genes that may contribute to larval growth and development.

This study provides a single-cell atlas of *E. granulosus* protoscoleces and identifies key candidate genes involved in larval development, providing a foundation for future functional studies and the exploration of potential anthelmintic targets.

## Introduction

Cystic echinococcosis (CE) is a zoonosis caused by infection with the metacestode stage of *E.granulosus* ^[1]^ and poses a substantial burden on human health and livestock production ^[2]^. CE occurs worldwide ^[3]^ and is particularly prevalent in parts of Africa ^[4]^, South America ^[5]^, the Mediterranean region^[6]^, Eastern Europe ^[7]^, and Central Asia ^[8, 9]^. The *E. granulosus* life cycle involves definitive hosts, primarily dogs, and intermediate hosts, including sheep and goats ^[9]^. Humans become accidental intermediate hosts through ingestion of parasite eggs. The eggs hatch in the digestive tract and release oncospheres, which penetrate the intestinal wall, reach the liver through the portal circulation, and develop into metacestodes that form cystic lesions ^[10]^.

The hydatid cyst comprises a cyst wall, cyst fluid, brood capsules, and protoscoleces. The cyst wall consists of an outer laminated layer ^[11]^ and an inner germinal layer ^[12]^. The germinal layer has a high nuclear density and proliferative activity and produces brood capsules toward the cyst interior, within which protoscoleces subsequently develop ^[13]^. As the cyst enlarges, it compresses surrounding tissues and is accompanied by local inflammation and fibrosis^[14]^. If untreated, cyst rupture and other complications may lead to organ failure or death^[9]^.

The capacity of the larva to grow and develop depends on stem cell populations capable of proliferation and differentiation. In flatworms, neoblasts contribute to tissue renewal, growth, and development in planarians and trematodes ^[15, 16]^. In cestodes, germinative cells are considered the only proliferative cells in the larval stage ^[17]^ and can give rise to differentiated cell types that support larval growth and development.

Drug treatment of CE currently relies primarily on benzimidazoles, which have limitations including poor absorption and hepatotoxicity ^[18, 19]^. There is therefore a need for safer and more effective anti-CE drugs. Understanding the cellular and molecular mechanisms underlying the developmental capacity of *E. granulosus* may help inform the development of improved treatments.

Previous bulk transcriptomic studies identified differentially expressed genes associated with the development of *E. granulosus* protoscoleces, providing insight into molecular changes ^[20, 21]^. However, bulk transcriptomics measures average expression across mixed cell populations and cannot readily resolve transcriptional differences between individual cell types.

Single-cell RNA sequencing allows the characterization of cellular composition and transcriptional heterogeneity at single-cell resolution, offering a strategy to investigate parasite development. Single-cell analysis of larvae of the related species *Echinococcus multilocularis* has revealed transcriptional heterogeneity among germinative cells ^[22]^. However, the cellular composition, germinative cell heterogeneity, and potential differentiation relationships in *E. granulosus* protoscoleces remain incompletely characterized.

Here, we used single-cell RNA sequencing to construct a transcriptomic atlas of *E. granulosus* protoscoleces, characterize their major cell types and potential lineage trajectories, and identify candidate developmental key genes. These findings provide a basis for investigating parasite development and potential therapeutic targets.

## Results

### Unsupervised clustering identified 13 distinct cell clusters

Approximately 200 protoscoleces were isolated from hydatid cysts in infected sheep livers and subjected to single-cell RNA sequencing (Fig. 1A). Single-cell transcriptomic libraries were prepared using the 10x Genomics Chromium platform and sequenced an Illumina platform, yielding 8,028 cells.After quality filtering, 7,612 cells were retained for downstream analysis. These cells had 300–3,000 detected genes, 300–8,000 unique molecular identifiers (UMIs), and a mitochondrial gene expression proportion of no more than 20% (Fig. 1B).Dimensionality reduction and visualization using uniform manifold approximation and projection (UMAP) revealed 13 cell clusters (Fig. 1C,D).

**Figure 1.**
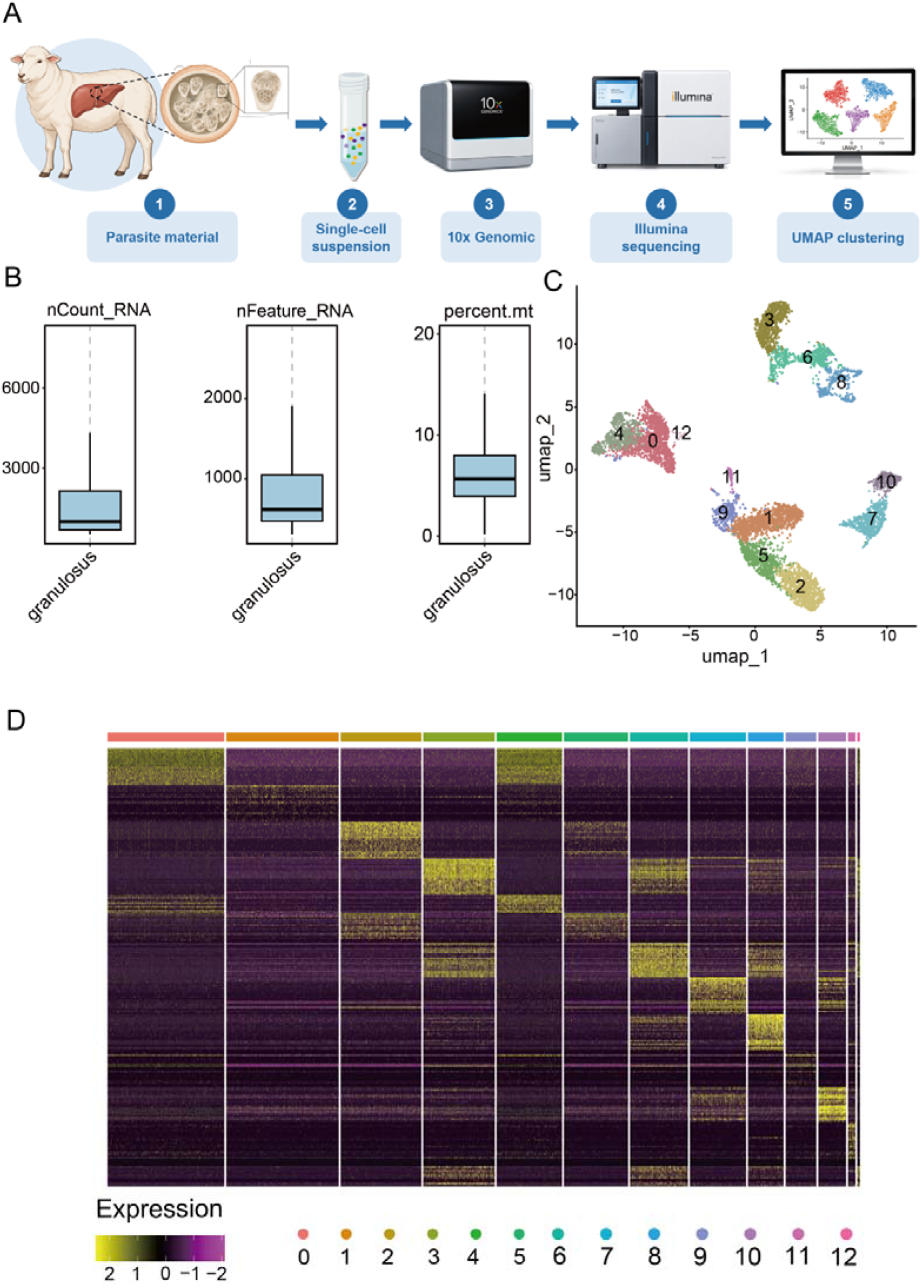
Single-cell transcriptome sequencing workflow and data preprocessing. (A) Overview of the experimental workflow. *E. granulosus* protoscoleces isolated from hydatid cysts in the livers of infected sheep were dissociated into single-cell suspensions. Single-cell RNA-sequencing libraries were prepared using the 10x Genomics Chromium platform and sequenced on an Illumina platform. Cell clusters with distinct transcriptional profiles were identified by unsupervised clustering and visualized using UMAP.(B) Boxplots showing the distributions of nCount_RNA, nFeature_RNA and percent.mt across all cells. (C) UMAP visualization of single-cell transcriptomes coloured by cell clusters (0–12). (D) Heatmap of top differentially expressed genes across clusters; colour represents normalized gene expression.

### Single-cell transcriptomics reveals six cell types with distinct transcriptional profiles

We further merged clusters with similar transcriptional profiles and annotated the resulting cell populations based on marker gene expression. For cell type annotation, we primarily referred to the markers reported by Loos et al. ^[22]^ in *E. multilocularis* larvae and mapped them to their one-to-one orthologs in *E. granulosus* (see Methods, Supplementary Data 1). For clusters without matching orthologous markers, cell identities were inferred using conserved molecular markers reported in other metazoans.

We identified six major cell populations (Fig. 2A,Supplementary Data 2): Germinative cells (1,934), Muscle (1,514), Neuron (1,172), Tegument (1,716), Storage (876), and Cilium (400). To further assess the annotations, we performed Gene Ontology (GO) Biological Process enrichment analysis of markers for each cell type (Fig. 2D). The enriched functions were consistent with the assigned cell identities(Supplementary Data 3).

**Figure 2.**
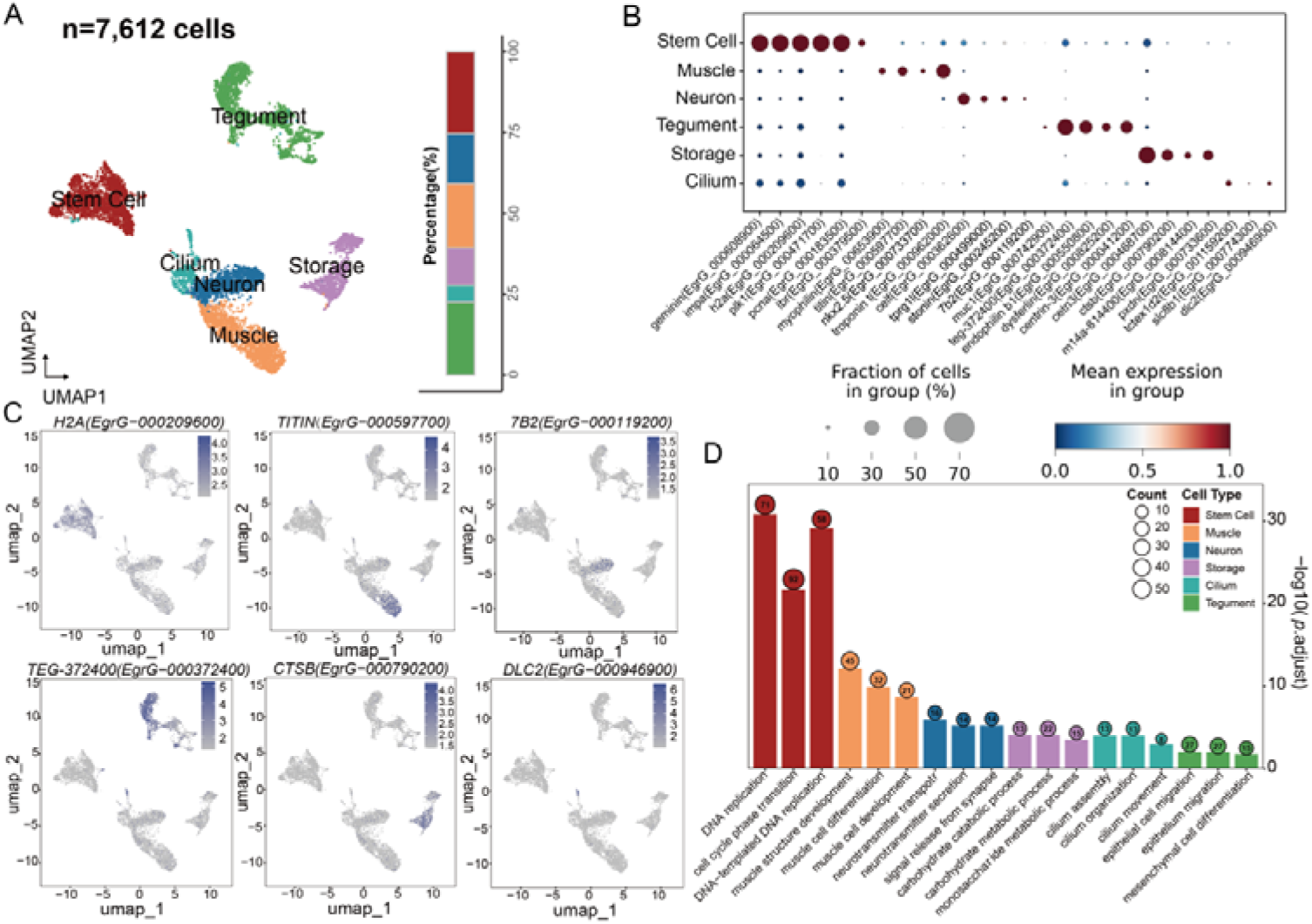
Single-cell transcriptomic atlas of the major cell populations in *E. granulosus* protoscoleces. (A)UMAP visualization of 7,612 single-cell transcriptomes from *E. granulosus* protoscoleces, classified into six major cell populations:Stem Cells,Muscle,Neuron,Tegument,Storage,and Cilium. The relative proportion of each cell population is shown below.(B)Dot plot displaying the expression of specific marker genes for each cell type in *E. granulosus*.(C)UMAP feature plots visualize the expression of cell-type-specific marker genes: H2A (Stem Cell), TITIN (Muscle), 7B2 (Neuron), TEG372400 (Tegument), CTSB (Storage), and DLC2 (Cilium). Colour intensity corresponds to the normalized gene expression level in individual cells.(D)Gene Ontology (GO) Biological Process enrichment analysis of marker genes for the major cell types. The x-axis shows representative GO Biological Process terms significantly enriched in each cell type, and the y-axis indicates enrichment significance [−log_10_(*P.adj*)]. Bar colors represent different cell types, while circle size and the numbers within the circles indicate the number of genes associated with each enriched term. The functional enrichment patterns of the different cell types were consistent with their respective annotations.

Germinative cells expressed representative genes including histone H2A (*h2a*; EgrG_000209600), proliferating cell nuclear antigen (*pcna*; EgrG_000183500), and *geminin* (EgrG_000608900) (Fig. 2 B,C). This population was enriched for biological processes related to DNA replication and the cell cycle, consistent with a proliferative state (Fig. 2D).

Muscle specifically expressed several canonical markers associated with muscle structure and contraction, including *titin* (EgrG_000597700), *myophilin* (EgrG_000653900), and *troponin T* (EgrG_000962000) (Fig. 2B,C). Terms associated with muscle development and differentiation were enriched in this population (Fig. 2D).

Neuron expressed genes including *tprg1l* (EgrG_000499000) and the neuroendocrine protein 7B2 (EgrG_000119200) (Fig. 2B,C). Enriched GO processes included neurotransmitter transport and secretion (Fig. 2D).

Tegument expressed *muc1* (EgrG_000742900), *teg-372400* (EgrG_000372400), *endophilin b1* (EgrG_000550800) and *annexin b18* (EgrG_000041200) (Fig. 2B,C). This population was enriched for processes including epithelial cell migration (Fig. 2D).

Storage expressed *ctsb* (EgrG_000790200), *m14a-814400* (EgrG_000814400), centrin 3 (*cetn3*; EgrG_000468700) and peroxidasin (*pxdn*; EgrG_000733600) (Fig. 2B,C). These cells were enriched for carbohydrate metabolic processes, consistent with their proposed metabolic function (Fig. 2D).

Cilium specifically expressed genes including the Tctex-type dynein light chain *tctex1d2* (EgrG_001159200) and dynein light chain 2 (*dlc2*; EgrG_000946900) (Fig. 2B,C). Terms related to cilium assembly and motility were enriched in this population (Fig. 2D).

Together, the transcriptional profiles and functional enrichment results provided complementary support for the cell type annotations.

### Transcriptional heterogeneity of germinative cells

Subclustering of germinative cells identified five subpopulations with distinct transcriptional profiles (Fig. 3A, B,Supplementary Data 4).

**Figure 3.**
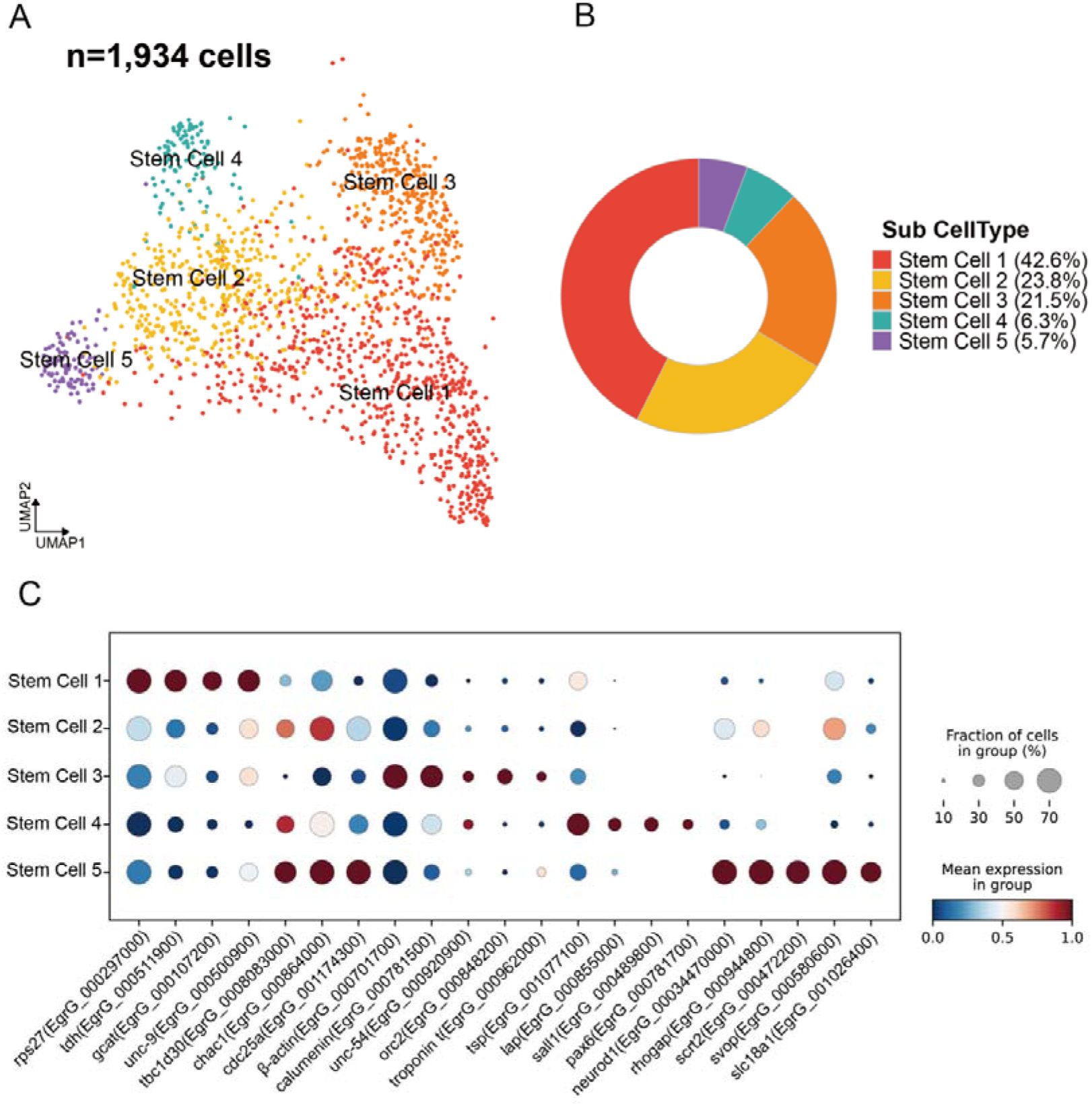
Identification and characterization of stem cell subpopulations. (A) UMAP visualization of 1,934 cells, coloured by five stem cell subpopulations (Stem Cell 1–5). (B) Donut chart showing the cell proportion of each stem cell subpopulations. (C) Dot plot illustrating the expression patterns of selected marker genes across the five stem cell subpopulations. The size of each dot represents the fraction of cells expressing the gene in the corresponding cluster, and the colour indicates the mean expression level.

Stem Cell 1 (n = 728) showed high expression of threonine dehydrogenase (*tdh*; EgrG_000511900), glycine C-acetyltransferase (*gcat*; EgrG_000107200), and the innexin gap junction protein unc-9 (EgrG_000500900) (Fig. 3C). No clear lineage-specific differentiation signal was detected in this subpopulation, suggesting an undifferentiated germinative cell state with a distinct transcriptional profile.

Stem Cell 3 (n = 367) retained stem cell-associated features while expressing muscle lineage-associated genes, including β*-actin* (EgrG_000701700), myosin heavy chain *unc-54* (EgrG_000092000), and *troponin T* (EgrG_000962000) (Fig. 3C). These findings suggest activation of a muscle lineage-associated transcriptional program and a possible muscle progenitor-like identity.

Stem Cell 4 (n = 108) highly expressed the tetraspanin *tsp* (EgrG_001077100) and leucine aminopeptidase *lap* (EgrG_000855500) (Fig. 3C). The expression of these genes, together with the previously reported tegument-associated germinative cell populations in *E. multilocularis* ^[22]^, suggests that Stem Cell 4 may represent a germinative cell population associated with the tegumental lineage.

Stem Cell 5 (n = 98) was enriched for genes associated with neurogenesis and neuronal differentiation, including *neurod1* (EgrG_000344700), rhogap (EgrG_000944800), and *scrt2* (EgrG_000472200) (Fig. 3C). The reported associations of neurod1 and rhogap with neural development in metazoans [22–25] support a possible early neural lineage state.

Stem Cell 2 (n = 407) lacked a prominent subcluster-specific expression signature but shared some transcriptional features with neural and tegumental lineage-associated subpopulations. It highly expressed the oxidative stress-associated gene *chac1* (EgrG_000864000) and the cell cycle regulator *cdc25a* (EgrG_001174300) (Fig. 3C). These features suggest a transitional state between undifferentiated germinative cells and lineage-associated progenitors.

These findings reveal transcriptional heterogeneity among germinative cells in *E.granulosus* protoscoleces, with distinct subpopulations displaying relatively undifferentiated, transitional, or lineage-associated transcriptional features.

### Potential differentiation trajectories and candidate developmental key genes

To investigate potential state transitions among protoscolex germinative cell subpopulations, we performed RNA velocity analysis. This approach uses the relative abundance of spliced and unspliced mRNA to estimate trends in transcriptional state changes and infer potential directions of differentiation (Fig. 4A).

**Figure 4.**
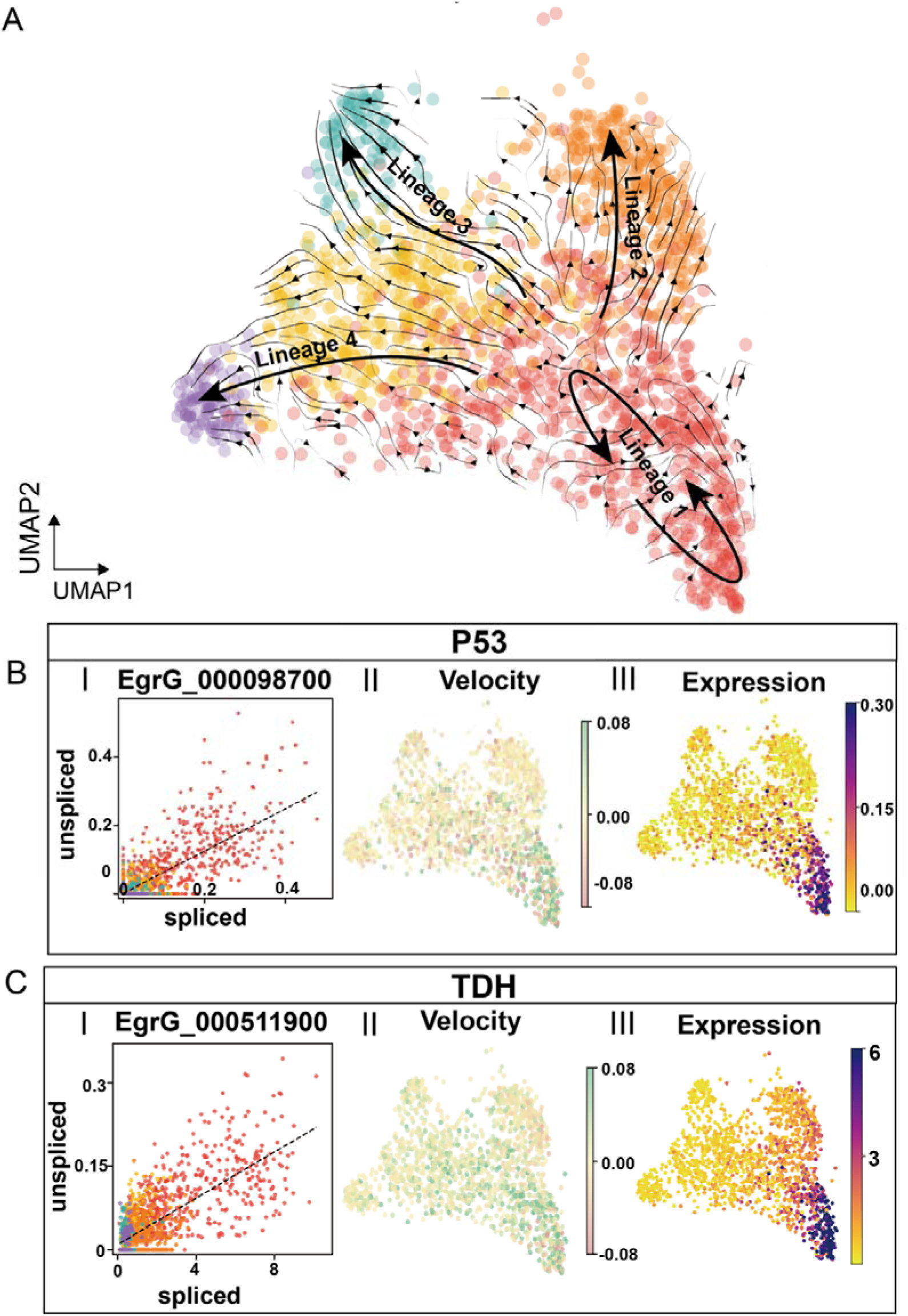
RNA velocity suggests potential differentiation trajectories and transcriptional dynamics of candidate developmental genes in *E. granulosus*. (A) UMAP embedding overlaid with RNA velocity stream field, showing four cell differentiation lineages (Lineage 1–4). Arrows indicate the inferred direction of cell state transitions.(B–C) Expression and RNA velocity of the P53 homolog (*EgrG_000098700*) and TDH (*EgrG_000511900*) across Stem Cell subpopulations. (I) Phase portraits showing the abundance of spliced and unspliced transcripts on the x- and y-axes, respectively. Dashed lines indicate model-fitted reference lines. (II) Gene-specific RNA velocity projected onto UMAP space, with colors indicating estimated velocity values. (III) Gene expression projected onto UMAP space, with colors indicating relative expression levels.

The analysis suggested four potential trajectories originating in Stem Cell 1, consistent with its relatively early cell state and absence of an obvious lineage-specific differentiation signature. Lineage 1 primarily involved transitions within Stem Cell 1 and may reflect processes associated with germinative cell self-renewal and population homeostasis.

Lineage 2 showed a trend from Stem Cell 1 toward the muscle progenitor-like Stem Cell 3 population. Lineages 3 and 4 suggested transitions from Stem Cell 1 through Stem Cell 2 toward the tegumental progenitor-like Stem Cell 4 and neural progenitor-like Stem Cell 5 populations, respectively. Stem Cell 2 may therefore represent an intermediate population in these transitions, consistent with its expression of some tegumental and neural lineage-associated markers.

To identify candidate factors potentially associated with germinative cell differentiation, we examined dynamic genes that fit the kinetic model together with markers of early germinative cells (see Methods). This analysis highlighted a *p53* homolog (EgrG_000098700) and threonine dehydrogenase (*tdh*; EgrG_000511900). Both genes showed velocity signals and high expression in Stem Cell 1(Fig 4B,C), suggesting active transcriptional programs in early germinative cells and a possible association with differentiation.

These analyses suggest potential differentiation trajectories of protoscolex germinative cells and identify candidate factors for future functional validation.

## Discussion

In this study, we generated a single-cell transcriptomic atlas of *E. granulosus* protoscoleces and characterized their cellular composition.

Based on transcriptional profiles, we identified six major cell populations: Germinative, Neuron, Muscle, Tegument, Storage, and Cilium. GO enrichment patterns were consistent with the proposed identities, providing additional support for the annotations.

Most of the cell types identified in our study were consistent with those reported in the single-cell transcriptomic atlas of *E.multilocularis* metacestodes constructed by Loos et al ^[22]^. The expression patterns of marker genes were highly conserved between the two species, suggesting that their corresponding cell types share conserved transcriptional features.We identified a cell population enriched for cilium movement-related processes that was not clearly identified in the previously reported *E. multilocularis* larvae atlas. We speculate that these cilium-associated cells may participate in physiological processes during the protoscolex stage.

Germinative cells are considered the only proliferative cells in *E. granulosus* larvae and underpin parasite growth and development ^[17]^. In addition to broadly expressed proliferation-associated markers such as *h2a*, geminin, and *pcna*, the lamin B receptor gene *lbr* was of interest. In many metazoans, *lbr* contributes to peripheral heterochromatin organization and changes in nuclear architecture associated with differentiation ^[26]^. Although its function in *E. granulosus* remains unclear, *lbr* may provide an entry point for investigating nuclear organization during germinative cell differentiation ^[22]^.

Subclustering identified five germinative cell subpopulations with distinct transcriptional profiles, indicating marked heterogeneity in protoscoleces.In our study, *lap*, *neurod1*, and *rhogap* discriminated germinative cell subclusters, as they did in the prior single-cell study of *E. multilocularis* ^[22]^,.pointing to shared transcriptional features of germinative cell subsets between the two species.However, *nos-1* (EgrG_000861500), *nos-2* (EgrG_000606200), *ago-2A* (EgrG_000739100), and *ago-2B* (EgrG_000911700), which have previously been used to examine germinative cell heterogeneity ^[17, 27]^, did not effectively distinguish protoscolex germinative cell subpopulations in our data. This finding may reflect differences in expression patterns between the two species.

The proposed intermediate state Stem cell 2 was implicated in transitions toward both neural and tegumental lineages, raising the possibility that these two lineages share the same early differentiation branch. This pattern resembles the ectoderm-associated differentiation routes reported in planarians by Zeng et al. ^[28]^. The separate muscle-associated branch involving Stem Cell 3 in Lineage 2 is also consistent with the differentiation of mesoderm-associated cells toward a muscle fate in planarians ^[28]^.This implies that the core framework of stem cell lineage differentiation exhibits partial evolutionary conservation among metazoans.

We identified a *p53* homolog and *tdh* as candidate developmental regulators. In *E. multilocularis*, Em-p53 is enriched in larval germinative cells ^[29]^ and has been implicated in oxidative stress responses and apoptosis ^[30]^. Recent studies have suggested that the role of invertebrate p53 family proteins extend beyond DNA damage responses and apoptosis and may include the regulation of stem cell proliferation, self-renewal, and lineage differentiation. For example, knockdown of *Smed-p53* in planarians causes abnormal stem cell proliferation and impaired differentiation^[31]^. In *E. multilocularis*, threonine dehydrogenase Em-TDH participates in L-threonine metabolism; supplementation with L-threonine promotes metacestode vesicle growth in vitro, and the enzyme has been proposed as a drug target for alveolar echinococcosis(AE) ^[32]^. Together with these previous findings, our results suggest that the *p53* homolog and *tdh* warrant investigation as candidate key genes of germinative cell differentiation and larval development in *E. granulosus*.

This study has limitations. Cell type annotations relied primarily on marker expression and functional enrichment and require spatial validation. The proposed germinative cell trajectories were mainly inferred from RNA-velocity algorithm and lack direct lineage-tracing evidence. In addition, the candidate developmental key genes were not experimentally tested; future studies using approaches such as in vitro gene perturbation will be needed to assess their functions.

In conclusion, this study characterizes the cellular composition of *E. granulosus* protoscoleces at single-cell resolution, identifies candidate markers for major cell types, reveals germinative cell heterogeneity, and proposes potential differentiation trajectories and candidate genes associated with larval development. These findings provide a foundation for further investigation of *E. granulosus* development and CE pathogenesis.

## Abbreviations

E. granulosus: Echinococcus granulosus
E. multilocularis: Echinococcus multilocularis

## Ethics approval and consent to participate

The animal study was reviewed and approved by the Ethics Committee of Inner Mongolia University. This research did not involve humans and/or human data, or material.

## Consent for publication

Not applicable.

## Acknowledgments

Not applicable.

## Author Contributors

QW conceived and designed the study. JWQ performed the experiments. QW and HYW conducted the data analysis. QW drafted the manuscript. All authors have read and approved the final version of the manuscript.

## Materials and Methods

### Reference genome construction

The *E. granulosus* nuclear reference genome and gene annotations were downloaded from WormBase ParaSite. Because the nuclear reference did not include mitochondrial sequences, the mitochondrial genome and its annotation were obtained separately and combined with the nuclear sequences and annotations to generate a reference containing both genomes for read alignment and gene expression quantification.

### 10x Genomics library preparation and sequencing

Individual cells were encapsulated with barcoded gel beads and reverse transcription reagents using the 10x Genomics platform to generate gel beads in emulsion (GEMs). Following synthesis, purification, and amplification of barcoded cDNA, libraries were prepared according to the Chromium Single Cell 3′ Reagent Kit v2 instructions (10x Genomics, USA). Libraries were sequenced on an Illumina HiSeq 4000 platform to generate 75-bp paired-end reads.

### Alignment and quality control of single-cell transcriptomic data

*E. granulosus* single-cell transcriptomic reads were aligned to the custom reference genome using the 10x Genomics Cell Ranger pipeline ^[33]^ (v2.2.0). Empty droplets were removed using the default Cell Ranger thresholds, and the filtered expression matrix was used for downstream clustering.

### One-to-one ortholog mapping

A homolog mapping table was generated based on protein sequence similarity ^[34]^. Protein sequences for the two species were downloaded and used to build local BLAST protein databases. Bidirectional BLASTp searches ^[34, 35]^ were performed to identify potential homologs. Hits were filtered by sequence similarity scores and statistical significance (*E-value* < 1 × 10), with reciprocal best hits preferentially retained to improve confidence in homolog assignment.

### Single-cell data analysis

Approximately 200 *E. granulosus* were pooled to prepare a single-cell suspension. Cells were captured and barcoded and transcriptomic libraries were generated using the 10x Genomics Chromium platform. Sequencing reads were aligned to the custom reference genome.

Quality control of gene expression count matrices was performed using Seurat^[36]^ package (v4.3.1). Putative doublets were subsequently identified and removed using DoubletFinder ^[37]^ (v2.0.4), with optimal parameters determined using the paramSweep function.

Data were normalized using NormalizeData and centered and scaled using ScaleData. For each dataset, 2,000 highly variable genes were selected using FindVariableFeatures, followed by principal component analysis (PCA). The first 15 principal components were used for neighbor graph construction, clustering, and uniform manifold approximation and projection (UMAP). FindNeighbors and FindClusters were used for clustering; 13 clusters were obtained for the *E. granulosus* dataset at resolution = 0.5. Pearson correlations between cluster-average expression profiles and hierarchical clustering were used to merge transcriptionally similar clusters, yielding six major cell populations.

### Protein functional annotation and enrichment analysis

*E. granulosus* protein sequences were submitted to the eggNOG-mapper online platform (v5.0) for functional annotation. The resulting data were processed in R and used to construct an *E. granulosus* GO annotation database with AnnotationForge (v1.44.0).

Differentially expressed genes between cell clusters were identified using Seurat FindMarkers with test.use = “wilcox”, only.pos = TRUE, min.pct = 0.25, and logfc.threshold = 0.25. Genes with *p_val* < 0.05 and *avg_log2FC* > 1 were selected for enrichment analysis. GO enrichment was performed using clusterProfiler ^[38]^ (v4.10.1) across the Biological Process (BP), Molecular Function (MF), and Cellular Component (CC) categories. Multiple testing was corrected using the Benjamini–Hochberg method, with FDR threshold of < 0.05.

### Germinative cell subclustering

To characterize heterogeneity within the germinative cell population, subclustering was performed using Seurat FindSubCluster with graph.name = “RNA_nn” and resolution = 0.5. Germinative cell subpopulations were then extracted using subset for downstream analysis.

### RNA velocity analysis

To further characterize the putative pluripotent stem cell populations, RNA velocity analysis was performed using the single-cell transcriptomic dataset of *E. granulosus* protoscoleces. Cell Ranger output was used to generate loom files containing spliced and unspliced transcript information with Velocyto ^[39]^ (v0.17.17). The dynamical model in scVelo (v0.2.2) was then used to model transient cell states and splicing kinetics and estimate velocity vectors for individual cells.

### Identification of dynamic genes

Candidate genes associated with cell state transitions were identified from the abundance of spliced and unspliced transcripts using the Velocyto function scv.tl.rank_velocity_genes withmin_corr = 0.5. Gene-specific RNA velocity and expression were visualized using scv.pl.velocity.

